# Molecular determinants of the modulation of the VSD-PD coupling mechanism of the K_V_7.1 channel by the KCNE1 ancillary subunits

**DOI:** 10.1101/2021.09.01.457404

**Authors:** Audrey Deyawe Kongmeneck, Marina A. Kasimova, Mounir Tarek

## Abstract

The IK_S_ current is diffused through the plasma membranes of cardiomyocytes during the last phase of the cardiac action potential. This repolarization current is conducted by a tetrameric protein complex derived from the co-expression of four voltage-gated potassium channel K_V_7.1 α-subunits and KCNE1 ancillary subunits from KCNQ1 and KCNE1 genes, respectively. We studied here the conformational space of K_V_7.1 in presence and absence of KCNE1, by building transmembrane models of their known Resting, Intermediate, and Activated states. We conducted Molecular Dynamics simulations of these models in lipid bilayers including the phosphatidyl-inositol-4,5-bisphosphate (PIP_2_) lipids. The comparative analysis of MD trajectories obtained for the K_V_7.1 and IK_S_ models reveals how KCNE1 shifts the coupling mechanism between the activation state of the Voltage Sensor Domain of the channel and the conformation (open or closed) of its Pore Domain.

## Introduction

IK_S_ channel is an oligomeric protein complex located in the plasma membrane of cardiomyocytes, among others. When a modification of transmembrane potential occurs, this channel opens and specifically diffuses potassium ions from the cytoplasm to the extracellular medium through the lipid bilayer. In cardiomyocytes, this voltage-gated potassium channel generates the IK_S_ current during the last fourth phase of the cardiac action potential (Barhanin et al., 1996; Nerbonne and Kass, 2005; Tristani-Firouzi and Sanguinetti, 1998). The structure of IK_S_ channel is a tetrameric protein complex derived from the co-expression of four K_V_7.1 α-subunits, along with a various number of β-subunits ranging from two to four (Nakajo et al., 2010). α-subunits and β-subunits result from the expression of KCNQ1 and KCNE1 genes, respectively.

As for most KV channels, K_V_7.1 α-subunits are organized to form a homotetramer whose voltage-sensor domain (VSD) and pore domain (PD) structures are swapped. Each α-subunit is composed of 6 transmembrane helical segments. The first four ones (S1-S4), whose fourth segment move across the membrane as a response to transmembrane voltage modification, form the VSD. The fourfold symmetrical structure of last two segments (S5, S6) forms the PD. The K_V_7.1 α-subunit also contains a cytoplasmic CTerm region composed of four cytosolic helices, the first two ones being connected to the sixth transmembrane segment (Sun and MacKinnon, 2017, 2020), the last two being located deeper in the cytosol, forming the tetramerization domain (Wiener et al., 2008). The primary sequence of the β-subunit KCNE1 counts 129 amino acids that are divided into one extracellular NTerm domain, one helical transmembrane (TMD) domain, and a cytosolic CTerm domain (Tian et al., 2007).

Over a hundred mutations of both KCNQ1 and KCNE1 genes have been reported to induce an inability of the protein complex to generate its IK_S_ outward current, in order to return cardiomyocytes membranes toward their resting potential. These loss-of-function mutations are associated with long QT syndromes (LQTS) (Kapplinger et al., 2009; Napolitano et al., 2005; Splawski et al., 2000) which are inherited diseases defined by an extended duration of the cardiac action potential. This delay in cardiac repolarization disturbs the propagation of the cardiac action potential throughout the myocardium tissue, leading to cardiac arrhythmias. Due to the localization of IK_S_ complexes in the inner ear (Nicolas et al., 2001), some KCNQ1 and KCNE1 mutations are also associated to the Jervell and Lange-Nielsen syndrome (JLNS), a particular inherited cardiac arrhythmia which also involves deafness (Jervell and Lange-Nielsen, 1957).

Several studies also reported the existence of single nucleotide polymorphisms (SNPs) in the KCNQ1 gene that are associated with the appearance of type 2 diabetes (Adeyemo et al., 2015; Al-Shammari et al., 2017; Li et al., 2017; Mussig et al., 2009). Hence, IK_S_ channel constitute a major therapeutic target for the treatment of these channelopathies (Kim, 2014). As a consequence, its function must be studied and well understood in order to be able to develop any potential drug (McGivern, 2007). This protein complex has been extensively studied over the last twenty years, through biophysical (Sun and MacKinnon, 2017, 2020; Taylor et al., 2020), physiological (Barro-Soria et al., 2014; Loussouarn et al., 2003; Osteen et al., 2010; Wang et al., 1999) and computational approaches (Eckey et al., 2014; Gofman et al., 2012; Jalily Hasani et al., 2017; Kang et al., 2008; Kuenze et al., 2020a; Xu et al., 2013; Zaydman et al., 2014).

As in many VSD of most voltage-gated ion channels (Bezanilla, 2000), the segment S2 in the VSD of the K_V_7.1 channel contains two evolutionarily conserved binding sites for S4 gating charges, the first one being E160 (E1), the second one including two acidic residues, E170 (E2) and D202 (D3), each located in S2 and S3 segments, respectively (Tao et al., 2010). This S4 conformational change, triggered by membrane depolarization, also involves a translation of these gating charges through an aromatic residue from S2, F167, which was suggested to separate the solvent accessible surface from the occluded site in the VSD. In addition, this residue is also conserved in the homologous Shaker KV channel (Lacroix and Bezanilla, 2011). Charge-reversal mutagenesis studies of the K_V_7.1 channel in the absence (Wu et al., 2010a) and the presence (Wu et al., 2010b; Zaydman et al., 2014) of KCNE1 allowed for the identification of the distinct stable states of the VSD of K_V_7.1, each defined by a specific set of salt bridges between S4 gating charges and their binding sites.

Numerous studies highlighted the biophysical properties of the IK_S_ channel complex as distinct from those of the K_V_7.1 channel: the ionic conductance is 4-fold increased, the voltage-dependence of activation is right-shifted towards depolarized voltages, and activation kinetics of the K^+^ current are slowed (Barhanin et al., 1996; Sanguinetti et al., 1996). Therefore, KCNE1 is suspected to modulate K_V_7.1 action by disrupting both its VSD activation and VSD-PD coupling mechanisms (Shamgar et al., 2008; Sun et al., 2012). Indeed, the pore opening mechanism of most KV channels is elicited by another mechanism called the VSD-PD coupling. The latter is defined as protein-protein interactions between VSD and PD of distinct α-subunits that aim to couple the activation state of the VSD (resting, Intermediate or activated) with the conformation (open or closed) of the PD (Roux, 2006). Charge reversal mutagenesis studies, among others, provided significant hints on the influence of KCNE1 subunits on the actual pore opening mechanism of the K_V_7.1 channel, as well as the VSD-PD coupling. The ionic currents obtained for each arrested state of the K_V_7.1 channel in absence of KCNE1, allowed one to predict if the VSD-PD coupling has occurred or not in each VSD state.

A recent study reported that in absence of KCNE1, the VSD-PD coupling of the K_V_7.1 channel occurs when the VSD is in both the intermediate and the activated state (Hou et al., 2017). The results obtained for K_V_7.1 in presence of KCNE1 shed light on the effects of KCNE1 on the ionic current of the K_V_7.1 channel in each of these three states: the current is reduced in the resting state, suppressed in the intermediate state, and enhanced in the activated state (Wu et al., 2010b; Zaydman et al., 2014).

Several studies were conducted in distinct KV channels in order to unravel the molecular determinants of the VSD-PD coupling mechanism. These studies shed light on the importance of specific interactions between the S4-S5L and the C-terminal region of S6, both located in the cytoplasmic side of the membrane (Choveau et al., 2011, 2012). Particularly, the cytoplasmic part of K_V_7.1 S6 segment was shown to be mainly involved in IK_S_ and K_V_7.1 pore gating (Boulet et al., 2007; Labro et al., 2011). The G/V curve of K_V_7.1, which represents its ionic conductance as a function of the transmembrane voltage, has a sigmoid shape, as encountered in most of the studied voltage dependent cation channels (Cui, 2016). This shape, along with the “domain-swapped” quaternary structure of K_V_7.1 subunits, suggests an allosteric mechanism for the activation of the voltage sensors of these channels, as well as for their VSD/PD coupling mechanism that leads to pore opening and ion conduction. Previous experimental studies showed that this allosteric coupling may be realized through a ligand/receptor mechanism in which S4-S5L acts an inhibitor, stabilizing the gate in a closed state (Labro et al., 2011). Upon membrane depolarization, S4 drags S4-S5L out of its S6 binding pocket, increasing the open probability of the channel. Despite of these significant insights of K_V_7.1 VSD-PD coupling mechanism, the molecular determinants allowing for the explanation of its modulation by KCNE1 subunits remains unresolved.

To that extent, tridimensional (3D) models of the IK_S_ channel in its open (Gofman et al., 2012; Xu et al., 2013) and closed (Kang et al., 2008) states have been constructed to provide an atomistic level knowledge about its structure and function. For these models, the location of KCNE1, between each K_V_7.1 subunit and facing both of their transmembrane domains in the IK_S_ complex, was determined by a series of various mutagenesis experiments (Chan et al., 2012; Chung et al., 2009; Lvov et al., 2010; Tapper and George, 2001; Wang et al., 2011; Xu et al., 2008, 2013). While these experimental studies highlighted several hints on the activation mechanism of the K_V_7.1 channel in presence of KCNE1, most of the IK_S_ 3D models that have been published for the last decade were modeled in order to investigate the molecular determinants of K_V_7.1-KCNE1 interaction. As a consequence, these computational results are not allowing for the prediction of a possible modulation mechanism of K_V_7.1 VSD- PD coupling by KCNE1. Recently, a 3D homology model (Jalily Hasani et al., 2017) of the IK_S_ complex was submitted to Molecular Dynamics (MD) simulations in a POPC:PIP2 membrane with a 10:1 ratio, but K_V_7.1-KCNE1 interactions were not extensively investigated in this study (Jalily Hasani et al., 2018). Hence, these models do not provide molecular comprehension of KCNE1 effects on the K_V_7.1’s VSD-PD coupling mechanism. While a host of studies suggests that IK_S_ complexes can be expressed with a 4:4 K_V_7.1:KCNE1 stoichiometry (Murray et al., 2016; Nakajo et al., 2010; Thompson et al., 2018; Wang et al., 2020) most of the aforementioned models have not been built with this ratio.

Recently, our integrative study of K_V_7.1 in absence of KCNE1 subunits suggested that the allosteric VSD-PD coupling mechanism of the K_V_7.1 channel is rather described by a “hand-and-elbow” gating mechanism (Hou et al., 2020). In this model, the S4 helix and S4-S5L form a bent arm. Upon membrane depolarization, the S4 segment (upper arm) moves upwards in two resolvable steps, first to the intermediate state and then to the activated state. Each step is described by a set of interactions between S4/S4-S5L region (elbow) and the S5 and S6 of a neighboring subunit. Both steps share a constant set of interactions between the C-terminus of S4-S5L (hand) and the cytoplasmic region of S6 (S6c) of the same subunit. Accordingly, the S4 (upper arm) movement toward the intermediate state triggers S4/S4-S5L region (elbow) upwards, while the C-terminus of S4-S5L (hand) pulls S6c in the pore to promote channel opening at both intermediate and activated states of the VSD.

The molecular determinants allowing for the modulation of K_V_7.1 function by KCNE1 are however yet to be investigated. To address these question, computational chemistry methods are insightful to unravel the elements of protein function at a molecular level of precision.

As several experimental studies reported the importance of the phosphatidyl-inositol-4,5- biphosphate (PIP2) on the gating of K_V_7.1 (Eckey et al., 2014; Loussouarn et al., 2003; Zaydman and Cui, 2014; Zaydman et al., 2013), we previously conducted an MD study (Kasimova et al., 2015a) of this channel’s open and closed states models. The resulting MD trajectories allowed one to localize the binding site of PIP2 interacting with K_V_7.1 subunits, and to characterize the key elements of K_V_7.1 modulation by PIP2 in absence of KCNE1. The lipid was shown to participate in VSD/PD coupling of the channel through state-dependent interactions, preventing repulsive forces between basic residues from S2-S3L and S4 in the resting/closed RC state of the channel , and between basic residues from S2-S3L and S6 in the AO state of the channel. The RC state refers to conformations of the channel where the VSDs are in the Resting state and the pore is in the Closed state, while the AO state refers to conformations of the channel where the VSDs are in the Activated states and the pore is Open.

These MD simulations suggested that PIP2 may also constitute a third binding site for S4 gating charges. Indeed, the lipid was found to form salt-bridges with residues R237 (R4) and R243 (R6) in RC model of the K_V_7.1 model, and not in the AO model (Deyawe Kongmeneck et al., 2021; Kasimova et al., 2015a).

In a previous MD study of IK_S_ models, we demonstrated that the presence of KCNE1 requires a second PIP2 binding site to satisfy the experimental constraints known for this complex (Deyawe Kongmeneck et al., 2021). Therefore, we built a system containing two PIP2 binding sites for IK_S_ models: the first PIP2 binding site, noted PIP2 intra, is located under the VSD, between S2-S3L and S4-S5L as seen in the models of K_V_7.1 channel alone (Kasimova et al., 2015a) ; the second binding site, PIP2 inter, is located between the CTerm regions of S6 and KCNE1. Each IK_S_ model was submitted to ∼500ns MD simulations, to obtain fully relax systems for each state of the channel. The resulting MD trajectories have been extensively analyzed and compared to the computational results obtained from our previous integrative study of K_V_7.1 channel. We used the structural differences between K_V_7.1 and IK_S_ models to predict the molecular determinants of K_V_7.1 VSD-PD coupling mechanism in presence of KCNE1 subunits. The MD trajectories obtained for IK_S_ models suggest that KCNE1 subunits are involved in a set of state-dependent interactions with the VSD-PD coupling interfaces we identified in K_V_7.1 models, both validated by functional studies. These interactions might modify the motion of S4-S5L unraveled by the “hand-and-elbow” model of VSD-PD coupling mechanism predicted for K_V_7.1 channel alone. The following part of this work presents the key interactions of this predicted mechanism.

## Results

The structural validation of the protein-lipid interactions in the IK_S_ models shed light on a set of three KCNE conserved basic residues. Located in the CTERM domain of KCNE1, these residues undergo a rotational movement during VSD activation, as described by its state-dependent electrostatic interactions with PIP2 in two distinct binding sites (Deyawe Kongmeneck et al., 2021). Here we investigated the hydrophobic interactions between KCNE1 and K_V_7.1. These interactions allowed us to predict how they could possibly induce the conformational changes in K_V_7.1 subunits that can explain the conductivity results obtained experimentally for each K_V_7.1 state, both in presence and absence of KCNE1 (Wu et al., 2010b, 2010a). To investigate the modulation of the K_V_7.1 VSD-PD coupling by KCNE1, we focused on the residues from the S4-S5L, S5, S6 and KCNE1 residues, and we monitored the distance to their closest residues (within a 5 Å radius).

As the presence of KCNE1 reduces drastically the ionic conductance of the K_V_7.1 channel in the Resting and Intermediate states, but increases it in the Activated state (Wu et al., 2010b, 2010a), we used this experimental data and our previous findings on the K_V_7.1 coupling mechanism (Hou et al., 2020) to investigate how KCNE1 interacts with K_V_7.1 subunits in the IK_S_ models to modify the K_V_7.1 channel VSD-PD’s coupling mechanism. To do this, we monitored specific state-independent intrasubunit interactions, as well as state-dependent intersubunit interactions, including those highlighted by our integrative study conducted on the K_V_7.1 models, in the MD trajectories we obtained for IK_S_ models.

The MD trajectories obtained for the K_V_7.1 models shed light on a cluster of residues in the pore domain, called hereafter the PD cluster. It is composed of 14 residues of the S4-S5L (W248 R249 L250 L251 V254 V255 F256 H258 R259), S5 (E261 L262) and S6 (I346 L347 V355) which overlap with the seventeen predicted molecular determinants of K_V_7.1 VSD-PD coupling in both IO and AO states (Hou et al., 2020). In IK_S_ models (Figure 1) and K_V_7.1 models, PD cluster residues are spread in two opposite surfaces (Figure S1), the first one being oriented towards the lipid bilayer (PD cluster outer surface), the second one being oriented towards the pore (PD cluster inner surface).

**Figure 1:**
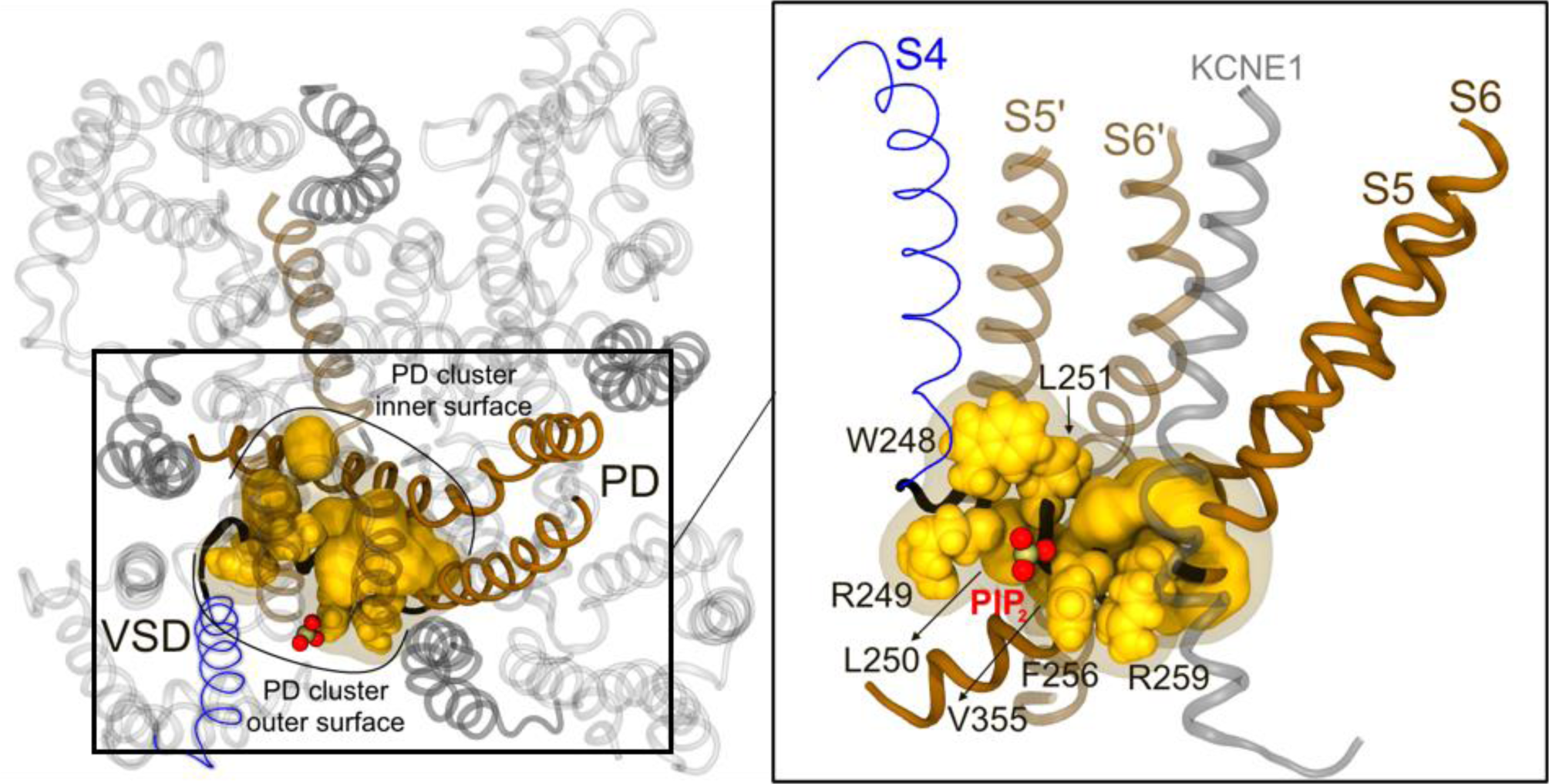
Structural mapping of PD cluster residues in IK_S_ AO state model. Cartoon representation of K_V_7.1 channel in its AO state in extracellular view (left) and the lateral view of the framed region (right). KCNE1 subunits are colored in dark grey. On both panels, PIP2 phosphate group is represented by red spheres. PD cluster outer surface residues are depicted in solid yellow spheres, while PD cluster inner surface residues are depicted in solid yellow surface. On the left panel, on one subunit, PD cluster residues are framed by a transparent yellow surface. S4 is colored in blue, S4-S5L in black, S5 and S6 in solid brown, while the segments S5’ and S6’ of the adjacent subunit are colored in transparent brown.

Here we assessed the interactions of PD cluster residues in the MD trajectories of K_V_7.1 and IK_S_ models. The results we obtained for K_V_7.1 models provided a thorough characterization of the molecular determinants of K_V_7.1 VSD-PD coupling mechanism. The results we obtained for IK_S_ models suggest that KCNE1 subunits induce a modification of the state-dependent interactions between PD cluster outer surface residues (W248 R249 L251 F256) and the rest of the channel (Figure S1). Those modifications appear to affect the interactions of PD cluster inner surface residues (V254 V255 H258 E261 L262 I346 L347 V355).

### Analyses of K_V_7.1 MD trajectories: The interactions between S4-S5L and S6 which mediate VSD-PD coupling mechanism are predominantly hydrophobic

In K_V_7.1 models, PD cluster inner surface counts a total of 10 residues that are engaged in two types of state-independent interactions: an electrostatic interaction between H258 and E261, along with VdW interactions between residues L250 V254 V255 H258, and residues V355 L347, L262 and I346, respectively (Figure S2). Among PD cluster outer surface residues (W248 R249 L251 F256), W248 and L251 are involved in state dependent interactions, including intersubunit ones with adjacent PD segments and intrasubunit ones with S4 segment.

Indeed, W248, bound to the bottom of adjacent S5 in the RC model, goes upwards along S5 to reach S4 in IO and AO models (Figure S3, upper panel); L251, bound to the bottom of adjacent S6 in the RC model, moves up along S6 to reach S5 in the IO and AO models (Figure S3, lower panel). Interestingly, the structural validation of K_V_7.1 MD trajectories suggests that PD cluster outer surface has three residues that are also involved in state independent interactions (Figure S2): The aromatic ring center of F256 is located close to the guanidinium carbon of R259 sidechain, allowing these residues to be engaged in a cation-π interaction, with frequencies higher than 0.6 in all models (Figure S2). R249 interacts with PIP2 with a frequency higher than 0.95 in every K_V_7.1 native state model through a salt-bridge interaction (Figure S1-S2).

The analysis of the MD trajectories obtained for K_V_7.1 models indicate that the motions of S4-S5L that underlie the VSD-PD coupling are driven by the remaining PD cluster outer surface residues, W248 and L251. Indeed, these residues are acting as hydrophobic hooks that allow PD cluster to be anchored in the PD of adjacent subunits as well as in the VSD of its own subunit, in a state dependent manner (Figure S4).

In the RC model (Figure S4, A), the benzene ring center of the W248 indole group is engaged in VdW interactions with the methyl group of T264 from S5 with a frequency of 0.75 (Figure S3, A). The terminal methyl groups of L251 are maintained between the sidechains of both L342 and I346 (Figure S3, B) from S6 via VdW interactions, with frequencies of 0.7 and 0.8, respectively.

In the IO model (Figure S4, B), the benzene ring center of W248 is interacting with the terminal methyl groups of L239 from S4 and I268 from an adjacent subunit (Figure S3, upper panel), while those of L251 are trapped between S5 and S6 segments through VdW interactions with I268 from S5 and L342 from S6 (Figure S3, lower panel). These suggests that during RC-IO transition, the S4-S5L has risen up along its adjacent S5 and S6 segments and has rotated anticlockwise towards its own VSD, allowing the linker to reach the bottom of S4. In the AO model (Figure S4, C), the sidechain of W248 is engaged in a set of VdW interactions between V241 from S4 and L271 from S5 of an adjacent subunit.

The other hook residue L251 still has its terminal methyl groups anchored between S5 and S6, but it has slightly translated upwards to engage in VdW interactions with F339 from S6 and I268 from S5. The interactions of both W248 and L251 in the K_V_7.1 AO model, which were validated experimentally in our previous integrative study (Hou et al., 2020), suggest that the PD cluster hook residues are facing S4 residues located deeper in the membrane during the IO-AO transition. The interactions in which L251 is engaged in K_V_7.1 AO state may be due to the upward movement of S4 segment during the IO-AO transition, during which the PD cluster has risen for a second time along its adjacent S5 and S6 segments while being still bound to S4 of its own subunit, leaving space for S6 cytoplasmic region to spread away from the pore axis toward the inner membrane surface (Figure S4, C).

Altogether, the state-dependent interactions of the PD cluster hook residues identified in K_V_7.1 models suggest that the S4-S5L undergoes subsequent upward translations (Figure S4, A-B, black arrows) and clockwise rotations (Figure S4, A-B, yellow arrows) of the PD cluster hook residues during the K_V_7.1 transitions from RC to IO model, then to the AO model. This mechanism appears to be induced by the motion of S4 upon the VSD’s activation, which is also characterized by subsequent upward translations normal to the bilayer, as well as subsequent clockwise rotations.

This S4 motion pulls the S4-S5L from the bottom of adjacent S5 and S6 segments in RC state to reach the middle of these segments and the bottom of S4 of its own subunit in the IO and AO states. Moreover, the intersubunit VdW interaction between L251 and I346 (Figure S3, lower panel) suggest that the latter might be crucial not only for the VSD-PD coupling, as shown in our previous study (Hou et al., 2020), but also for the stabilization of the relative position of S4-S5L with respect to S6c when the VSD is in its resting state (Figure S4, A). This relative position might be crucial for the initiation of K_V_7.1 VSD-PD coupling upon resting- intermediate VSD transition. As a consequence, the loss of the I346-L251 interaction induced by the I346A mutation might lead to a loss of function of K_V_7.1, characterized by an inability of VSD activation in response to membrane depolarization (for I346A).

Overall, these results confirm the molecular determinants of the VSD-PD coupling we previously validated for K_V_7.1 channel, in which S4-S5L motion drags the cytoplasmic region of S6 helices during both RC-IO and IO-AO transitions of K_V_7.1, and therefore induce the spreading of its S6 helices away from the pore axis during pore opening.

### KCNE1 subunits disturb the VSD-PD coupling mechanism of K_V_7.1

The MD trajectories of the IK_S_ models suggest that KCNE1 induce a modification of state-dependent interactions between the PD cluster outer surface residues and the rest of the channel. To that extent, we conducted additional analyses to identify among K_V_7.1-KCNE1 interactions those which might explain how KCNE1 subunits can modify these PD cluster interactions.

In our last reported study in which we investigated the interactions between KCNE1 subunits and PIP2 lipids in IK_S_ complex through computational methods, we shed light on the fact that KCNE1 undergoes a rotational movement during VSD activation, as described by its state-dependent electrostatic interactions with PIP2 in two distinct binding sites (Deyawe Kongmeneck et al., 2021). Here we investigated the hydrophobic interactions between KCNE1 residues and those of K_V_7.1.

From the IK_S_ trajectories, the state independent interactions in which PD cluster outer surface residues R249 F256 and R259 are involved in K_V_7.1 models are impaired: R249 still interacts with PIP2 (intra) through a salt-bridge interaction, with a frequency higher than 0.8 in the IK_S_ RC and IC models (Figure 2). However, this interaction frequency drops at 0.44 in the IK_S_ AO model which indicates that this interaction is either unstable or present in less than three subunits out of four throughout the MD trajectory. The aromatic ring center of F256 is not close enough to the guanidium group of R259, preventing these residues from engaging in a cation-π interaction in all IK_S_ models.

**Figure 2:**
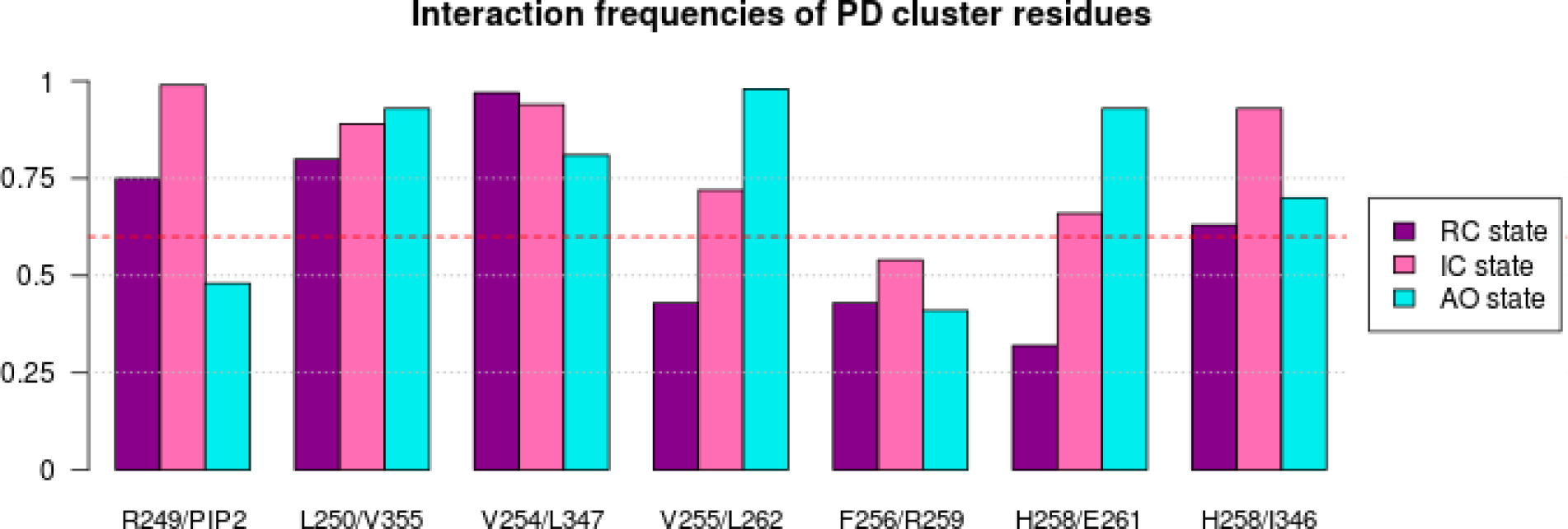
Intrasubunit interactions of PD cluster residues in IK_S_ models. The barplots display the frequencies of the state-dependent intrasubunit interactions involving PD cluster residues throughout the MD trajectories obtained for IK_S_ RC (purple), IC (pink), and AO (cyan) models. The frequency threshold of 0.6 is marked by a red dashed line.

The PD cluster hook residues W248 and L251 behave differently in the IK_S_ models (Figure 3, A). In the RC state, W248 is trapped between T264 and I268 with frequencies of 0.6, indicating that W248 is less buried in the membrane in the IK_S_ RC model compared to the K_V_7.1 RC model (Figure 4A, bottom). L251 interactions remain unchanged in the IK_S_ RC model with respect to K_V_7.1, as its methyl groups are bound to those of L342 and I346 with frequencies of 0.8 and 0.95, respectively (Figure 3, B). These interactions between the hook residues and the adjacent PD segments show that, in the RC state of K_V_7.1, the PD cluster residues are slightly less buried in the membrane in the presence of KCNE1.

**Figure 3:**
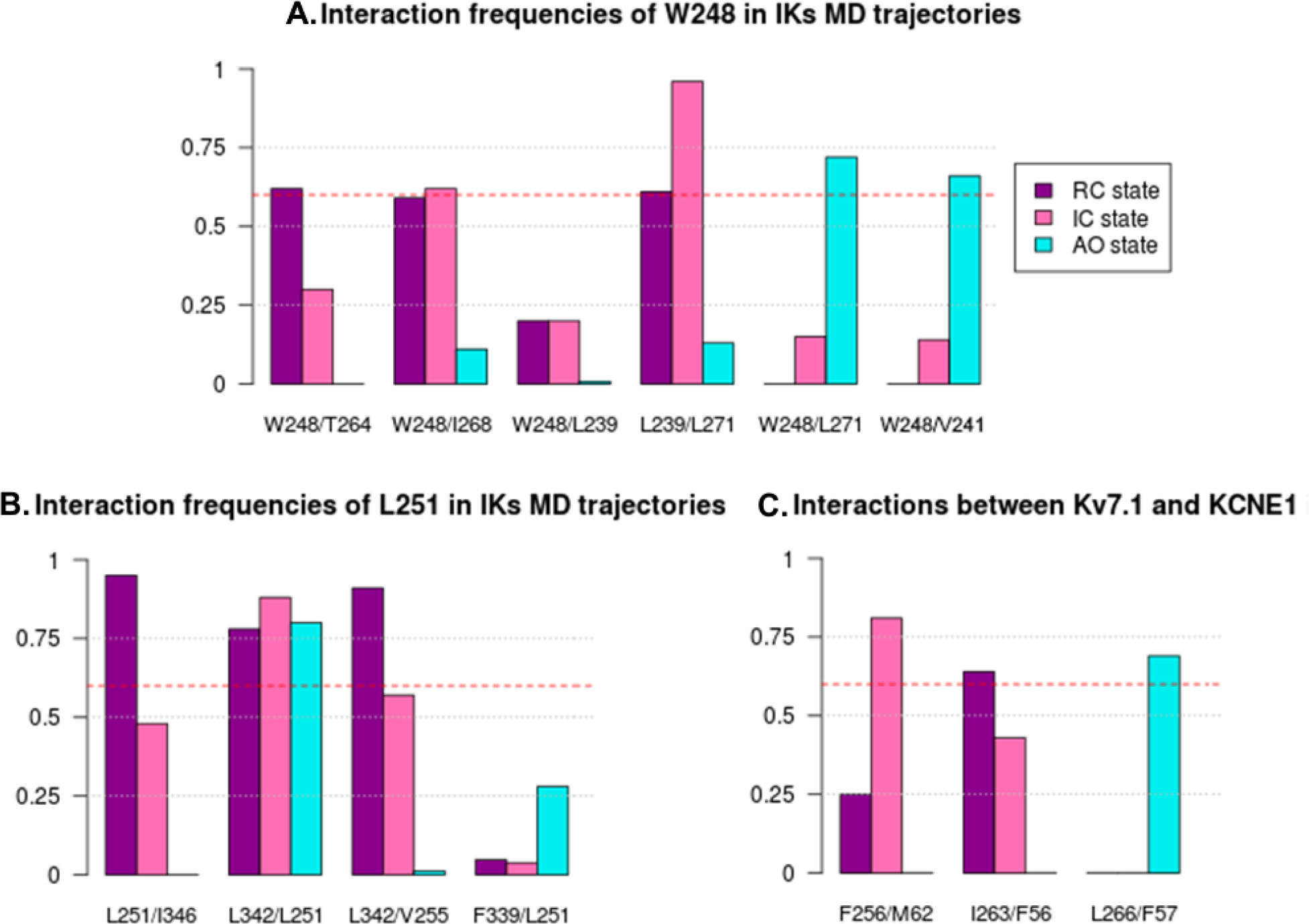
Intersubunit interactions of PD cluster residues are modified in IK_S_ models. The barplots display the frequencies of the state-dependent intersubunit interactions involving PD cluster outer surface residues and KCNE1 residues throughout the MD trajectories obtained for IK_S_ RC (purple), IC (pink), and AO (cyan) models. The frequency threshold of 0.6 is marked by a red dashed line.

**Figure 4:**
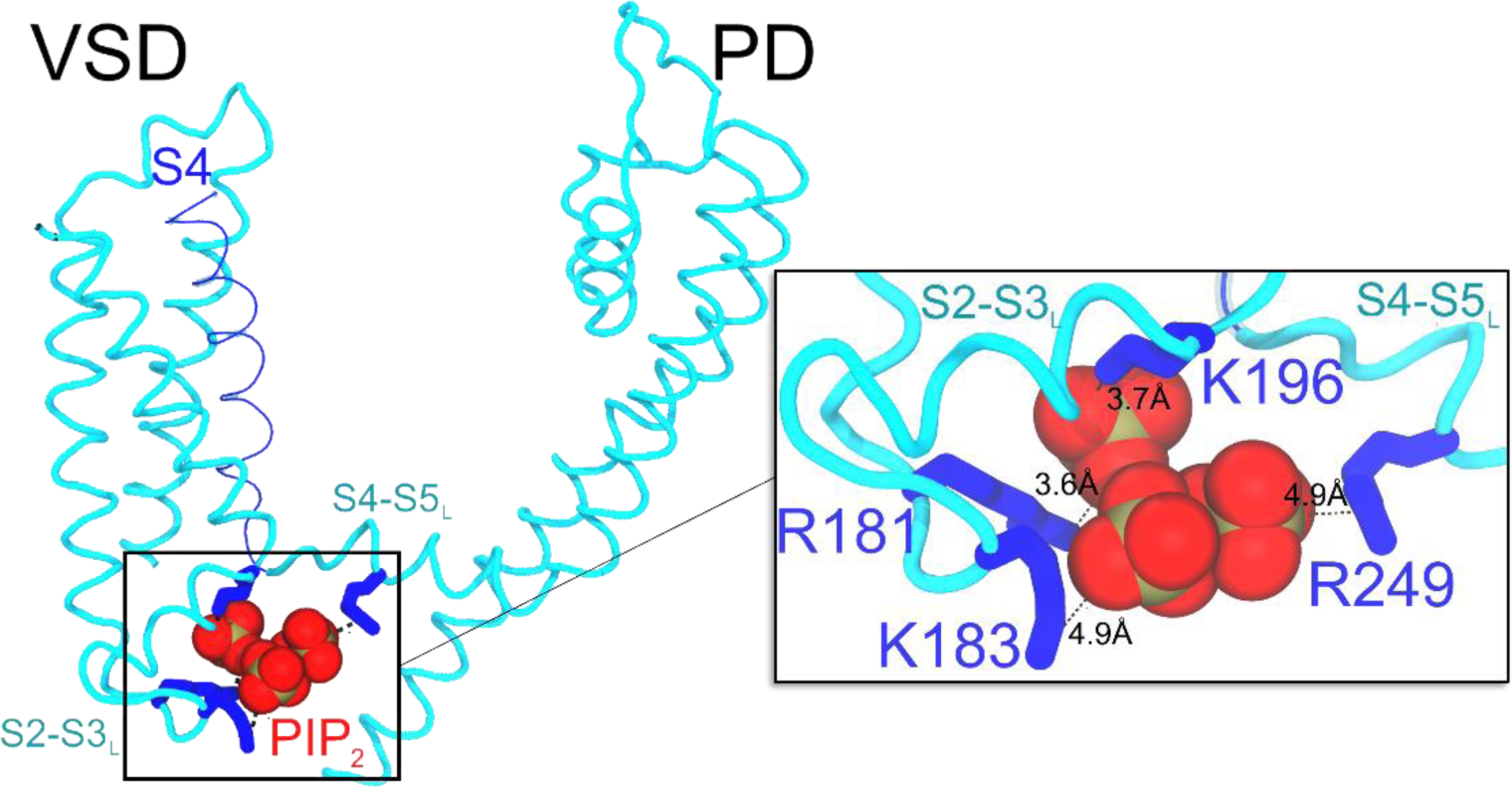
Position of PIP2 in the CryoEM structure of K_V_7.1:KCNE3 complex. The left panel shows the location of PIP2 phosphate groups (in red sphères) in the structure of K_V_7.1:KCNE3 complex (PDB ID: 6v01). The transmembrane region of K_V_7.1 subunit is shown in cyan cartoon, except S4 in transparent blue ribbon, and the closest basic residues to PIP2 (in blue sticks), located in S2-S3L (R181, K183, K196) and S4-S5L (R249). The right panel shows an extended view of the phospholipid, along with the specific distances (in Å) between its phosphate groups and the charge moieties of its closest K_V_7.1 basic residues. KCNE3 subunit is hidden for clarity.

In IK_S_ IC model, PD cluster hook residues are not binding S4, as the interaction frequency between W248 and L239 has dropped to 0.2. This interaction is impaired since W248 sidechains are located two far away from L239 ones in IC model (Figure 4, B). Instead, W248 aromatic ring remains bound to I268 methyl groups, at a frequency of 0.6. L251 sidechain leaves I346 to bind I268 as well, with a frequency of 0.86 (Figure 3, B). In AO model, W248 has finally reached S4, as its aromatic ring center is engaged in VdW interactions with methyl groups of both V241 from S4 and L271 from S5, with frequencies of 0.6 and 0.7, respectively (Figure 3, C). Meanwhile, L251 side chains remain bound to both I268 from S5 and L342 from S6, with frequencies of 1 and 0.8, respectively. In addition, the aromatic ring center of residue F256 from the outer surface of the PD cluster is engaged in a Met-Arom interaction with the sulfur atom of M62 from KCNE1, with a frequency of 0.8 (Figure 3C). The energy associated with Met-Arom interactions are known to be in the same range as salt-bridges interactions (Valley et al., 2012), so one can predict that this interaction might disrupt the motion of S4-S5L during RC-IC transition.

In the IK_S_ AO model, the K_V_7.1 AO state specific interactions are mostly present, except for the interaction between L251 and F339 (Figure 3, C). Instead, L251 still binds L342 with a frequency of 0.8 (Figure 3, B). This discrepancy may be due to the additional interaction between the KCNE1 TMD and the middle region of S5. Indeed, the VdW interactions between F54 from KCNE1 and L266 from S5 is present at a frequency of 0.7 (Figure 3, C). This interaction might pull the S4-S5L/S5 region away from the pore axis, and farther from its adjacent S6 segments. Furthermore, the PD cluster inner surface is slightly modified (Figure 1, left panel). The terminal methyl groups of L250 sidechains interact with both the aromatic ring center of F351 and the methyl groups of V355, with frequencies of 0.9 and 0.75, respectively (data not shown). Since F351 is located above V355 along S6 segment and, considering the relative position of S4-S5L with respect to the membrane in K_V_7.1 activated state, this interaction indicates that the cytoplasmic region of S6 segments is closer to the inner membrane surface in presence of KCNE1.

Analyses of PD cluster residues interactions in our IK_S_ models also show that state-independent interactions involving inner surface residues are slightly modified in RC and AO models with respect to K_V_7.1 models (Figure 4, A, C). Indeed, in RC model, S5 residues E261 and L262 from S5 are not interacting with the rest of PD cluster residues. As a result, V255 is bound to L342 from an adjacent S6 segment instead of L262 (Figure 3, A), with a frequency of 0.91 (Figure 3, B). Moreover, the aromatic ring center of residue F56 from KCNE1 TMD interacts through a VdW interaction with residue I263 from S5 segment at a frequency of 0.64 (Figure 3, C). Considering the position of I263, next to the PD cluster inner surface residues E261 and L262, one can assume this additional interaction with KCNE1 might modify the shape of PD cluster, leading it to be engaged in an intersubunit interaction with an adjacent S6.

These observed modifications with respect to K_V_7.1 models in IK_S_ RC state, in which S5 residues are split off from the PD cluster, may allow for a deeper burying of S4-S5 linker in the bilayer, and thereby for a narrower pore, as shown in by pore radii calculations of IK_S_ RC models (Deyawe Kongmeneck et al., 2021).

In IK_S_ IC model (Figure 4, B), PD cluster inner surface is finally reassembled, as S4-S5L may have undergone an upward movement upon RC-IC transition of K_V_7.1 subunits. As a consequence, S5 residues E261 and L262 are bound to the inner surface of PD cluster (Figure 2), but V255 still interacts with adjacent L342 with a frequency of 0.6 (Figure 3, B). This interaction might prevent the PD cluster to reach S4 as in K_V_7.1 IO model, as it remains bound to an adjacent subunit in the presence of KCNE1.

In IK_S_ AO model, PD cluster inner surface is still conserved. The terminal methyl groups of L250 sidechains interact the aromatic ring center of F351 from S6, with a frequency of 0.8. Since F351 is located above V355 along S6 and considering the relative position of S4-S5L with respect to the membrane in K_V_7.1 activated state, this interaction indicates that S6 segments are closer to the membrane in presence of KCNE1. In AO model, L251 does not interact with F339, as seen in K_V_7.1 AO model. Instead, L251 remains bound to L342 with frequency of 0.8 (Figure 3, B). This discrepancy may be due to additional interactions between KCNE1 TMD and the middle region of S5 (Figure 3, C). Indeed, VdW interactions between F54 from KCNE1 and F270 from S5, as well as between F57 from KCNE1 and L266 from S5, are both present at a frequency of 0.7. The presence of these interactions in AO models is probably due to PIP2. Located in the inner leaflet of the POPC bilayer, the lipid forms a bridge between S2-S3L and S4-S5L, which ensures VSD-PD coupling in both K_V_7.1 and IK_S_ AO states (Deyawe Kongmeneck et al., 2021; Kasimova et al., 2015b). Interestingly, in the CryoEM model of K_V_7.1-KCNE3 complex, the PIP2 lipid is located in the same area as in our MD systems (Figure 4).

Altogether, the K_V_7.1-KCNE1 state-dependent interactions we found in our IK_S_ models suggest consecutive clockwise rotations of the ancillary subunit during VSD activation of IK_S_ channel (Figure 5, grey arrows). In addition, these interactions induce a modification of state dependent interactions involving PD cluster outer surface residues. In IK_S_ RC model, the presence of KCNE1 turned out to stabilize the position of PD cluster residues along its adjacent PD segments (Figure 5, A). IK_S_ IC model, the presence of KCNE1 appeared to prevent the translation of S4-S5L from its adjacent PD segments towards S4 segment of its own subunit (Figure 5, B), which may prevent pore opening as well, as shown in model pore radii values we previously reported (Deyawe Kongmeneck et al., 2021). In IK_S_ AO model, S4- S5L appears to be eventually anchored to the bottom of S4, while KCNE1 binds the middle of S5 segment, as well as S6 cytoplasmic region to stabilize the open state of the PD (Figure 5, C). These interactions indicate that KCNE1 may pull S5 segments away from the pore axis, in the same direction of S4-S5L translational motion upon VSD activation (Figure 6, grey arrows).

**Figure 5:**
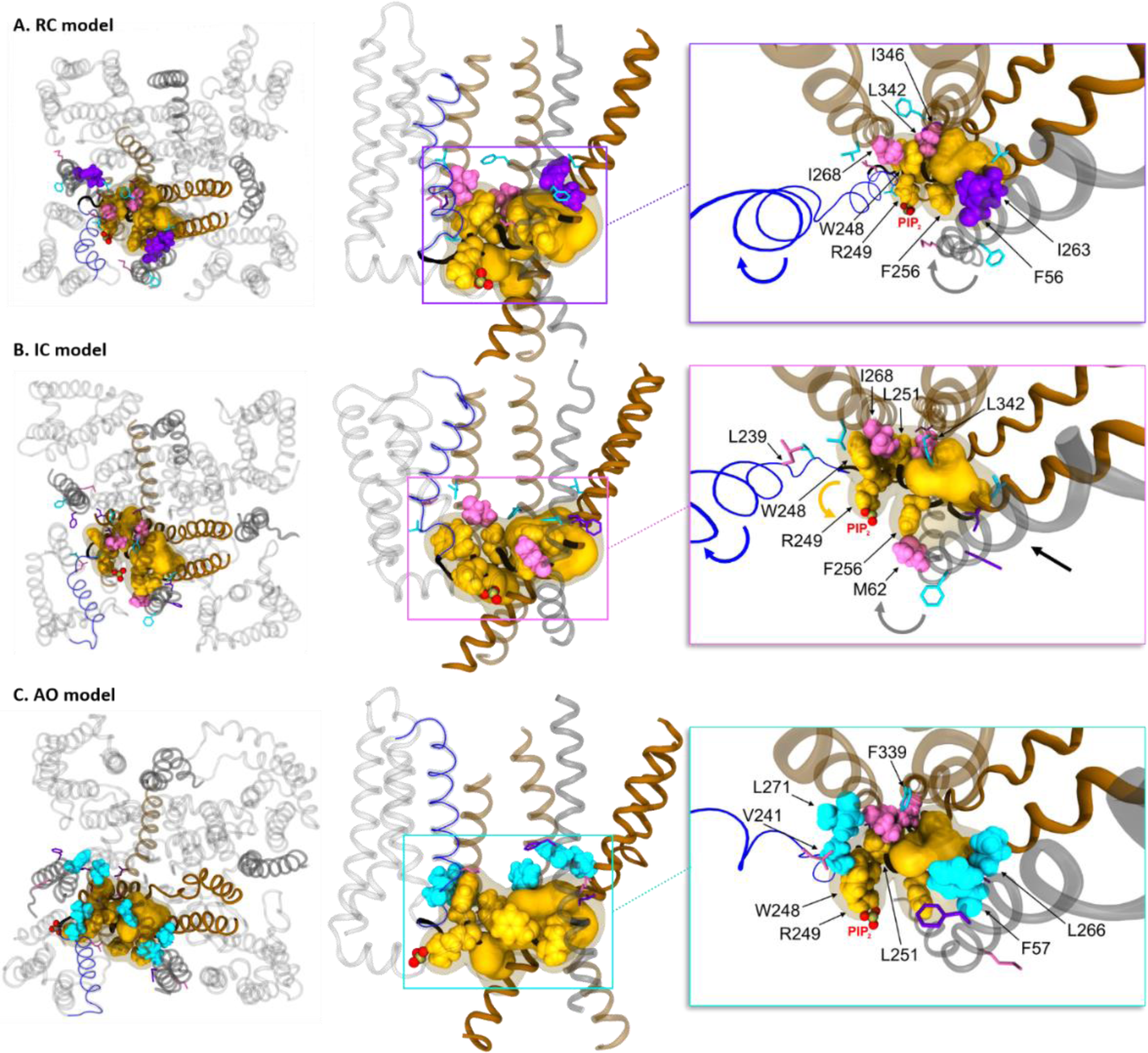
State dependent VSD-PD coupling interfaces in IK_S_ models. Each panel shows a cartoon representation of PD cluster residues in top view (left), lateral view (middle) and zoomed view (right) of **A.** RC model, **B.** IC model and **C.** AO model of IKS. In all panels, S4 is colored in blue, S4-S5L in black, S5 and S6 in solid brown, while the segments S5’ and S6’ of the adjacent subunit are colored in transparent brown. The entire PD cluster is depicted in transparent yellow surface, and PIP2 phosphate groups in red spheres. Hook residues W248, R249, L251 and PD cluster inner surface residues are represented in solid yellow spheres and surface, respectively. In each panel, residues which interact with PD cluster are represented in spheres, while those which do not are represented in sticks. The residues which interact with PD cluster in **A.** RC models are purple-colored, while those which interact in **B.** IO model are pink-colored, and those which interact in **C.** AO models are cyan-colored. Rotational motions of S4, and PD cluster residues W248, R249, L251 are represented by blue, and yellow arrows, respectively. Translational movement of PD cluster is represented by a black arrow.

**Figure 6:**
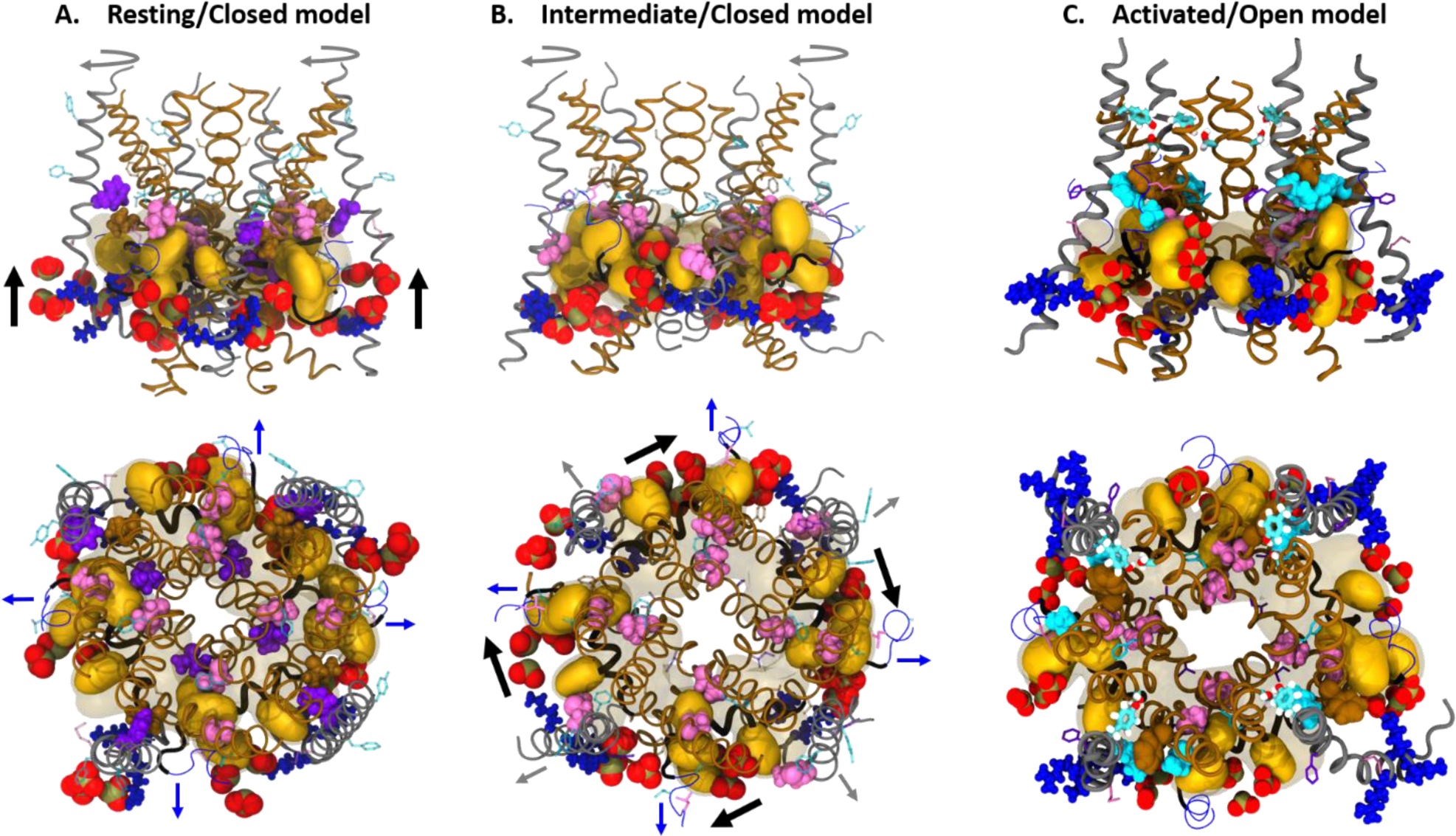
Prediction of K_V_7.1 coupling mechanism which mediates RC→IC and IC→AO transitions of PD cluster in presence of KCNE1. Side view (top) and extracellular view (bottom) of IK_S_ transmembrane segments (in ribbons) involved in VSD- PD coupling mechanism, such as S4 (blue), S4-S5LINKER (black), PD segments (brown) and KCNE1 (gray). PD cluster residues are represented in transparent yellow surface, while PD cluster hook residues W248 R249 and L251 are represented in solid yellow surface. Residues that interact with PD cluster are represented in spheres, while those which do not are represented in transparent sticks. These residues are colored in purple in **A.** RC model, in pink in **B.** IO model and in cyan in **C.** AO model, in which residues S330 and Y46 from S6 and KCNE1, respectively, are depicted in thicker cyan sticks. KCNE1 basic residues and PIP2 phosphate groups are depicted in blue and red spheres, respectively. Translational motions of S4, S4-S5LINKER and KCNE1 are represented by blue, black, and grey arrows, respectively.

Consequently, we proposed a model for the modulation mechanism of K_V_7.1 VSD-PD coupling by KCNE1 subunits, in which the ancillary subunits act as Velcro strips in the “hand-and-elbow” model (Hou et al., 2020) of the VSD-PD coupling mechanism of K_V_7.1 (Figure 7).

i. During the VSD’s activation from the resting to the intermediate state, the rotation of KCNE1 places its lower Velcro strip (located on the KCNE1 M62 residue) in front of F256, located on the outer surface of the K_V_7.1 PD cluster, blocking the rotation of the elbow and the hand, as well as its translation along the inner membrane surface that promote pore opening.
ii. During the VSD transition into the activated state, the second clockwise rotation of KCNE1 releases the lower Velcro strip, allowing for the elbow and the hand to rotate counterclockwise to reach both S4c and adjacent PD segments and promote channel opening. The rotation of KCNE1 places another Velcro strip (located on KCNE1 residue F57) in front of L266 on S5. These strips are assumed to pull PD segments farther from the pore axis than in K_V_7.1 AO state, which can explain the pore radii differences observed between K_V_7.1 and IK_S_ AO models (Deyawe Kongmeneck et al., 2021).

**Figure 7:**
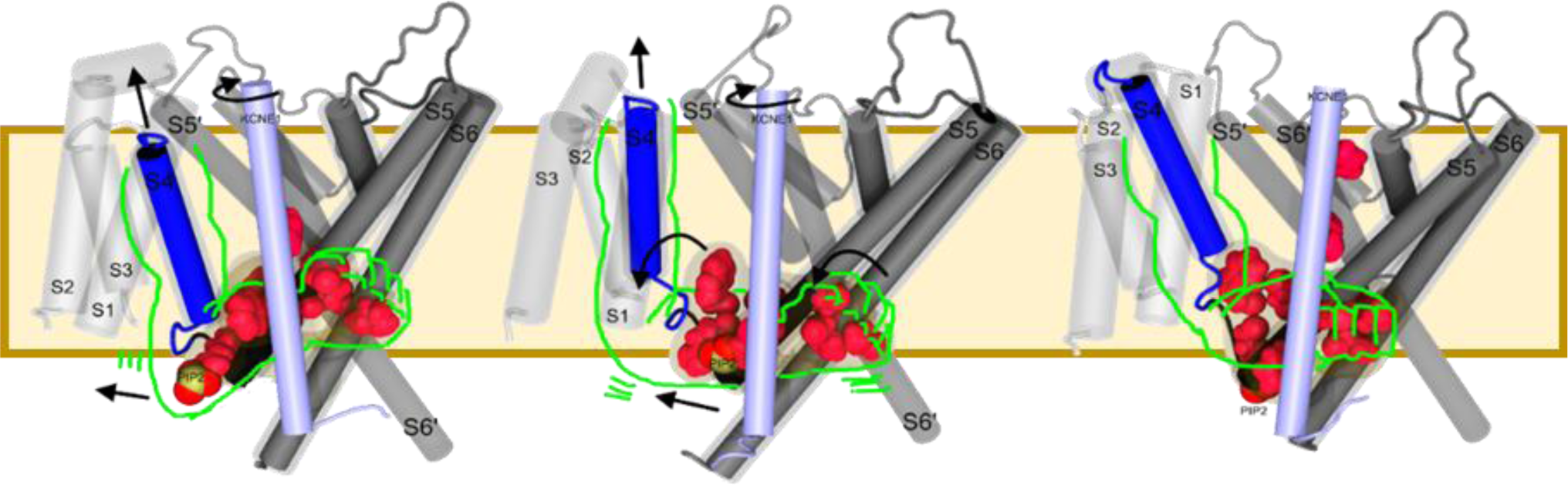
Hand-and-elbow model for IK_S_ state-dependent VSD-PD coupling mechanism. This figure shows a cartoon representation of K_V_7.1 transmembrane segments, including one entire subunit on the foreground with the VSD (transparent light gray) S4 (blue), S4-S5L (black), PD segments (solid gray) and KCNE1 segments (light blue) along with the PD segments of an adjacent subunit (transparent gray). The “elbow” residues W248 R249 L251, the “hand” residues H258 L262 I346 L347 and KCNE1 residues 46 57 62 are depicted in pink spheres. PIP2 phosphate groups in state independent interaction with the elbow (R249) are shown in red spheres. The entire arm that models the whole conformational changes is drawn in green. The motions of both the elbow and the arm during RC-IO and IO-AO transitions are depicted by black arrows.

Interestingly, a recent computational study, which consisted in the use of machine learning methods to generate a conformational space of IK_S_ channels, have yielded to the design of a structure-based predictor of IK_S_ channel experimental properties including its subconductance and gating current (Ramasubramanian and Rudy, 2018). The two sequential translations and rotations of S4 and the rotations of KCNE1 leading to VSD activation we predicted from our models are supported by the results obtained with this structure-based predictor. The fact that the molecular determinants of the mechanism of KCNE1 modulation has been unraveled by two distinct computational studies involving different methods is quite promising. Moreover, the rotational motion of KCNE1 subunits during the VSD’s activation was also suggested by previous functional studies (Chung et al., 2009; Xu et al., 2008). Altogether, these results increase the trustworthiness of our predictions.

## Discussion

When the concept of allostery is applied to voltage-gated ion channels, pore opening/closing transition is considered as the main function of the allosteric protein (Changeux, 2012). In addition, the VSD activation and deactivation occur when S4 moves along the electric field created by membrane depolarization and hyperpolarization, respectively. Accordingly, the VSD of K_V_7.1 can adopt three distinct conformations that induce PD opening/closing which itself directly depends on the VSD-PD coupling. This mechanism, considered here as an allosteric one according to the Monod-Wyman-Changeux (MWC) criteria, takes place at the interface between S4-S5L and the cytoplasmic part of S6. As such, this state-dependent S4- S5L/S6 interface is considered here as an allosteric site. In a protein which satisfies the MWC kinetic model, its function is defined by conformational changes that involve concerted motions at the level of this allosteric sites.

In the frame of the allosteric properties of the IK_S_ complex, the state-dependent VSD-PD coupling interfaces we identified for the K_V_7.1 IO state and both the K_V_7.1 and IK_S_ AO states are indeed inducing a sort of catalytic effect on the channel which is characterized by the opening of the pore to conduct potassium ions. KCNE1 acts as an antagonist in both resting and intermediate states of K_V_7.1, inhibiting the VSD-PD coupling mechanism by disrupting its key state-dependent and state-independent interactions (Figures 5-7). In the activated state, KCNE1 acts as an agonist of the coupling due to a conformational change which allows the ancillary subunit to engage in protein-protein interactions which may probably stabilize the open conformation of the pore, as well as the activated conformation of the VSD (Figures 3, 5C). This might explain how the IK_S_ current is sustained for a larger duration than the current generated by K_V_7.1 subunits in absence of KCNE1. The allosteric aspect of K_V_7.1 channel modulation by KCNE1 unraveled in this present study agrees with the results obtained from a recent integrative study in which several KCNE1 residues identified in this study (Y46, F57) were shown to have a significant impact on the mechanism that allows KCNE1 to slow K_V_7.1 activation in IK_S_ channel complex (Kuenze et al., 2020b). In addition, their computational study reveals that K_V_7.1 channel modulation by KCNE1 also involves proximities between S5 residues (I263, L266, Y267 and F270) and KCNE1 FTL motif residues (56 FTL 59), which also agrees with the state-dependent interactions we identified in this current study (Figure 5C, 6C).

The set of exquisite results we collected from MD simulations of our models of K_V_7.1 channel in presence of KCNE1 allowed us to propose a description of two simple movements which explain its VSD-PD coupling mechanism. This mechanism is characterized by a motion of the PD cluster, which translates upwards from the bottom of an adjacent S6, while rotating towards the bottom of S4 of its own subunit (Figures 6-7). Indeed, in K_V_7.1 models (Figure S4) the PD cluster residues are spread in two opposite areas with specified missions. The outer pore surface contains the hydrophobic residues W248 and L251 which are engaged in state-dependent interactions, depicting an upward movement of S4-S5L along its adjoining PD segments to reach the bottom of S4 and S2-S3L through R249-PIP2 interaction in the AO model.

This motion has been shown to be triggered by the VSD activation, defined by an upward motion of S4 along the electric field induced by membrane depolarization. The inner pore surface contains the S4-S5L residues engaged in state-independent VdW interactions with both S5 and S6 residues from the same subunit, which suggests a pulling of S6 segments from the pore axis toward the inner membrane surface upon S4 and S4-S5L movements during the VSD activation. The specificity in our IK_S_ models in relation with the various IK_S_ models reported so far (Eckey et al., 2014; Jalily Hasani et al., 2017, 2018; Kang et al., 2008; Kuenze et al., 2020b; Ramasubramanian and Rudy, 2018; Xu and Rudy, 2018) resides in the fact that the presence of PIP2 is acting as a hook to ensure the leverage effect of S4-S5L on S6 cytoplasmic region. Indeed, R249 from S4-S5L is interacting with PIP2 intra in all models, while PIP2 intra itself progressively anchors S2-S3L during S4 activation, while maintaining the S4-S5L close to the inner membrane surface.

Interestingly, the absence of PIP2 in the KCNQ1EM structure results in a S4-S5L located far from the S2-S3L and rather bound to the S6 cytoplasmic region, leading to the three major constriction zones which close its conduction pathway.

Specifically, the MD analyses conducted on our IK_S_ models has shown that the presence of KCNE1 modifies the VSD-PD coupling mechanism of K_V_7.1 channel in such a way to split the movements of PD cluster hook residues into two distinct transitions (Figure 6-7). The upward translation of the PD cluster, driven by L251, occurs during the IK_S_ RC-IC transition, while the anticlockwise rotation of the PD cluster, driven by W248, occurs during the IK_S_ IC- AO transition. At the end of the RC-IC transition, the IK_S_ channel is more likely to be closed, as the VSD-PD coupling mechanism is not completed in presence of KCNE1. Indeed, the protein-lipid component of this mechanism constitutes a tourniquet for S6 helices, preventing them from reaching the inner membrane surface, and leading to the interaction between the PD cluster and S4 as seen in both K_V_7.1 IO and AO models.

Besides, the presence of KCNE1 involves the participation of a third hook residue, F256. Located on the outer surface of the PD cluster, it directly interacts with KCNE1 in the IK_S_ IC model, which may hinder the progression of S4-S5L along the membrane inner surface when the VSD adopts its intermediate conformation. As a consequence, the conduction pathway of the IK_S_ IC state is more likely to be closed, as found in its pore radii calculations (Deyawe Kongmeneck et al., 2021). During the IC-AO transition, KCNE1 loses its interaction with PIP2 intra, which is allowed to anchor the PD cluster to the bottom of S4 segment and S2-S3L in the AO state, while shifting the S6 cytoplasmic region away from the pore axis to open the pore (Figure 6, C). Through an additional H-bond interaction between Y46 and S6 residue S330 in the extracellular side of the lipid bilayer (Figure 6, C, cyan sticks) KCNE1 may enhance the ionic current of K_V_7.1 subunits. As a matter of fact, this interaction may possibly explain the experimental results obtained for the IK_S_ channel mutant Y46A (Gofman et al., 2012), whose current amplitude is lower than the wild-type IK_S_ channel current. These results have provided a molecular scale comprehension of the numerous experimental studies which unraveled the key aspects of VSD-PD coupling and the way KCNE1 can possibly modulate this mechanism.

Overall, we have shown that protein-protein and protein-lipid components of K_V_7.1 VSD-PD coupling mechanism are mediated by a cluster of 12 residues which are mostly hydrophobic and involved in state-dependent intersubunit VdW interactions between S4-S5L and both S5 and S6 segments, as well as state-independent intrasubunit VdW interactions between hydrophobic residues from S4-S5L and S6. This coupling mechanism we unveiled is similar to the mechanical lever model, in which S4 upward movement drags a S4-S5L stuck to S6 cytoplasmic region, leading to pore opening of homologous KV1 channels (Choveau et al., 2012), as well as the reported integrative study conducted on homologous Shaker channel (Fernández-Mariño et al., 2018). The large number of VdW interactions we have determined for the PD cluster residues testify for their importance in the coupling mechanism. Indeed, among the 12 PD cluster residues, 10 are associated with LQTS when deleted or mutated into residues whose sidechains are of either different size or chemical property (Tab 1). For aliphatic residues, these pieces of information provide additional evidence of the importance of VdW interactions in conformational changes undergone by IK_S_ ion channels during membrane depolarization. Altogether, these innovative results make our 3D models reliable and robust enough to be used for more advanced computational studies of the function of IK_S_ complex, as their features allow for the explanation of the macroscopic currents measured on K_V_7.1 channel *in vitro* in presence and absence of KCNE1 and/or PIP2 lipids (Wang et al., 2020). The extended dataset of conformations generated by the MD simulations of our models of K_V_7.1 can be used to perform further investigations of the mechanistic aspects we predicted. For instance, free-energy calculations would allow for the determination of the energetic aspects of the transitions between the stable states of K_V_7.1 subunits, in absence and presence of KCNE1. Such calculations have already been conducted on the homologous KV1.2 channel (Delemotte et al., 2015, 2017), and the obtained results are shown to provide crucial information in the comprehension of its VSD activation.

**Table 1:** PD cluster residues associated with LQTS mutations.

| Kv7.1 residue | Segment | LQTS mutation(s) (reference) |
| --- | --- | --- |
| W248 | S4-S5L | W248R (Franqueza et al., 1999) |
| L250 |  | L250H (Itoh et al., 1998) L250P (Kapplinger et al., 2009) |
| L251 |  | L251P (Deschênes et al., 2003) |
| V254 |  | V254M (Choi et al., 2004; Donger et al., 1997; Wang et al., 1996) V254L (Tester et al., 2005) |
| V255 |  | Deletion V254-V255-F256 (Tester et al., 2005) |
| F256 |  |  |
| H258 |  | H258R, H258N (Tester et al., 2005) |
| R259 |  | R259H (Millat et al., 2006) R259L (Choi et al., 2004)<br>R259C (Jongbloed et al., 2002; Kubota et al., 2000; Van Langen et al., 2003) |
| E261 | S5 | E261K (Donger et al., 1997)<br>E261Q, E261D (Kapplinger et al., 2009) |
| L262 |  | L262V (Tester et al., 2005) |

In addition, brownian dynamics simulations can also be performed on these models in order to estimate the state-specific conductances of K_V_7.1 channel *in silico* [375, 376]. The results obtained would provide additional weight to the robustness and the reliability of our models in the prediction of the molecular determinants of K_V_7.1 channel function. In addition, the mechanistic aspects we determined can be used to study the interactions of K_V_7.1 channel (Bendahhou et al., 2005) and IK_S_ complex (Manderfield and George, 2008; Toyoda et al., 2006) with other KCNE ancillary subunits, as they were reported to modulate their function. In the case of IK_S_ complex modulation, the investigation can be facilitated since new CryoEM structures of K_V_7.1 subunits were resolved along with both PIP2 and KCNE3 ancillary subunits (Sun and MacKinnon, 2020). Our mechanistic results can as well be used to unravel the function of related KV channels, as the key residues we identified and validated through functional studies turned out to be evolutionarily related with most domain-swapped KV channels (Hou et al., 2020) through statistical coupling analysis (Rivoire et al., 2016).

## Material and Methods

To investigate the structural determinants of the VSD-PD coupling mechanism of the K_V_7.1 channel in presence of KCNE1, we first needed to build molecular models which had to be as trustworthy as possible with respect to experimental data available. Since no experimental structure of the K_V_7.1 channel was available yet, we used homology modeling to build our K_V_7.1 models (Deyawe et al., 2018). As the activation mechanism of K_V_7.1 involves three stable states. We modeled each state to obtain a larger spectrum of possible α-subunits conformations, allowing for the prediction of possible transition mechanisms for IK_S_ channel. For these models, only residues 122 to 366 from KCNQ1 human sequence, corresponding to the transmembrane region of the channel, were considered. To adjust the salt-bridge patterns of the VSD with respect to experimental results (Wu et al., 2010b, 2010a), and thereby to obtain distinct activation state models of the VSD, the charged group of E160 (E1) was constrained towards R237 (R4) in activated/open (AO) model, using the refined crystallographic structure of KV1.2 channel as a template (Chen et al., 2010); towards R231 (R2) in Intermediate/Closed (IC) model using the γ conformation of KV1.2 refined structure obtained from previous MD simulations (Delemotte et al., 2011) and towards R228 (R1) in Resting/Closed (RC) model, using the ε conformation of KV1.2 refined structure obtained from the aforementioned study.

The alignment of K_V_7.1 human sequence and KV1.2 rat sequence was conducted automatically, using ClustalW2 and refined manually and locally. For each state, 50 K_V_7.1 models were generated along with the TMD (residues 39 to 76) of human KCNE1 NMR structure (Tian et al., 2007), using a K_V_7.1:KCNE1 subunit ratio of 4:4 using MODELLER (Eswar et al., 2007).

Among these 50 obtained models, the ten best ranked models according to the knowledge-based scoring function DOPE (<u>D</u>iscrete <u>O</u>ptimizing <u>P</u>rotein <u>E</u>nergy) (Shen and Sali, 2006) were selected for stereochemical quality evaluation using PROCHECK (Laskowski et al., 1993) software.

For Molecular Dynamics (MD) simulations, we used the input generator of CHARMM (Jo et al., 2008), to add Palmitoyl-Oleyl PC (POPC) molecules around the protein structure (Wu et al., 2014) and a PIP2 molecule at the bottom of each VSD (PIP2 intra) of KCNE1 subunits (PIP2 inter) with respect to experimental studies (Eckey et al., 2014; Li et al., 2011; Zaydman et al., 2013)and our previous computational results of MD simulations conducted on K_V_7.1 subunits along with PIP2. The systems (lipid+channel) were surrounded by two slabs of 150 mM [KCl] solution. Simulations were carried out using the CHARMM36 force-field (Huang and Mackerell, 2013), along with the CMAP correction (Buck et al., 2006). The MD systems were simulated in a NPT ensemble for the equilibration of the systems to reproduce experimental conditions, at 300K and 1 atm, with a time-step of 2.0 fs, using NAMD code (Phillips et al., 2005) to perform all MD calculations.

As we assessed the reliability of MD trajectories obtained for both K_V_7.1 (Hou et al., 2020) and IK_S_ models (Deyawe Kongmeneck et al., 2021) with respect to experimental data in previous computational studies, in this present work we only investigated the modulation of K_V_7.1 VSD-PD coupling by KCNE1, by focusing on the interactions of residues from S4-S5L, S5, S6 and KCNE1. To do that, we monitored the distance with their closest residues (within a 5 Å radius).

We considered only interhelical contacts and, according to the physicochemical properties of the considered side chain, the neighbor residue search was limited to those with similar chemical properties. For example, to look for the neighbor residues of an aromatic one, we restrained the search on hydrophobic residues.

Moreover, depending on the nature of the concerned residue, we computed distance calculations to characterize several types of interactions that are usually found in transmembrane protein structures, using the same cut-off values as in our previous MD study (Hou et al., 2020). For each interatomic distance, we calculated the frequency at which the latter is under the threshold values mentioned above in all subunits throughout the MD trajectory of each model. We considered the interaction as present in all the subunits of the model if this frequency is above 0.6. All these analyses were performed with the use of several home-made script programs written in TCL language and executed within VMD shell (Dalke and Schulten, 1997; Humphrey et al., 1996).

## Supplementary Material

**Figure S1:**
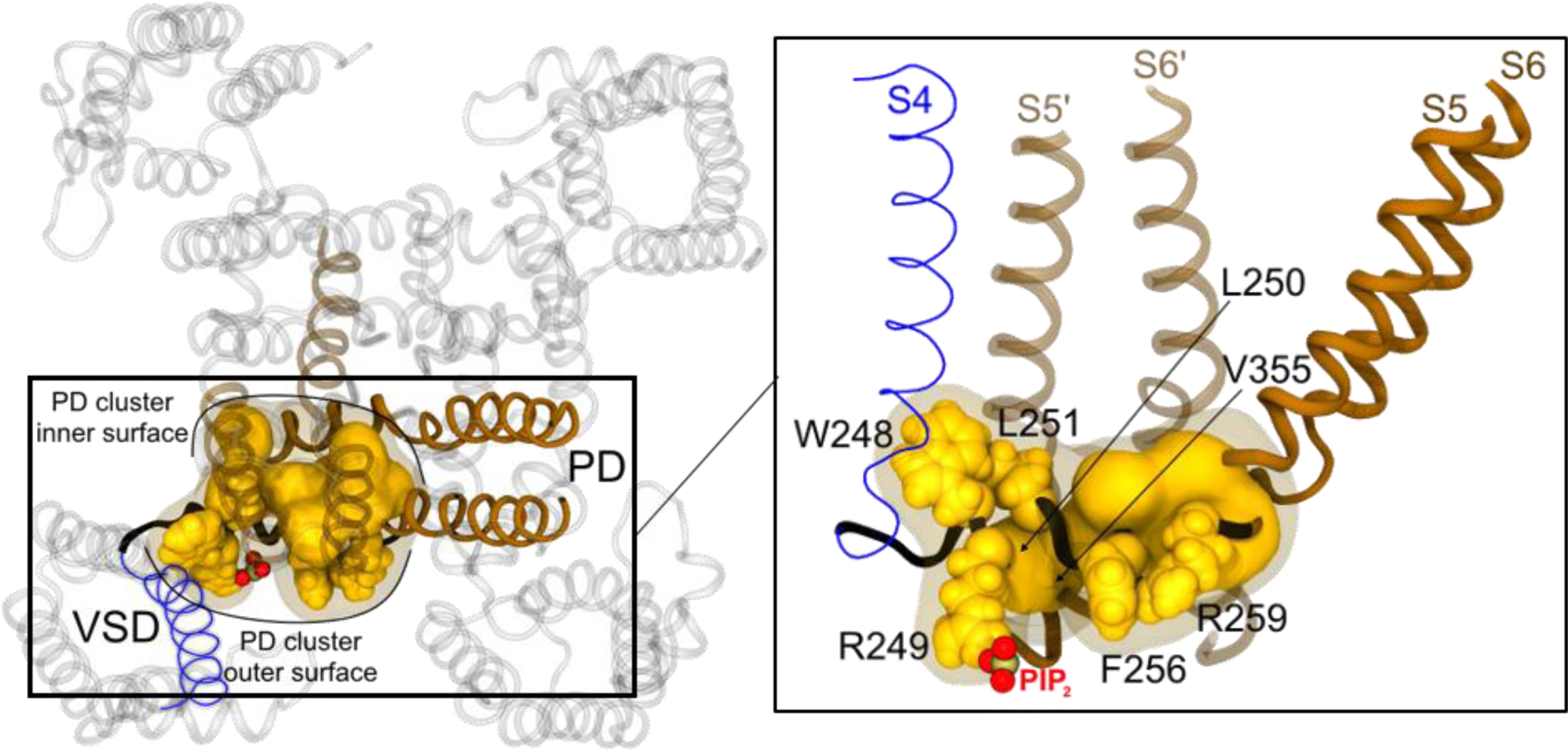
Structural mapping of PD cluster residues in K_V_7.1 3D model. Cartoon representation of K_V_7.1 channel in its AO state in extracellular view (left) and the lateral view of the framed region (right). On both panels, PIP2 phosphate group is represented by red spheres. PD cluster outer surface residues are depicted in solid yellow spheres, while PD cluster inner surface residues are depicted in solid yellow surface. On the left panel, on one subunit, PD cluster residues are framed by a transparent yellow surface. S4 is colored in blue, S4-S5L in black, S5 and S6 in solid brown, while the segments S5’ and S6’ of the adjacent subunit are colored in transparent brown.

**Figure S2:**
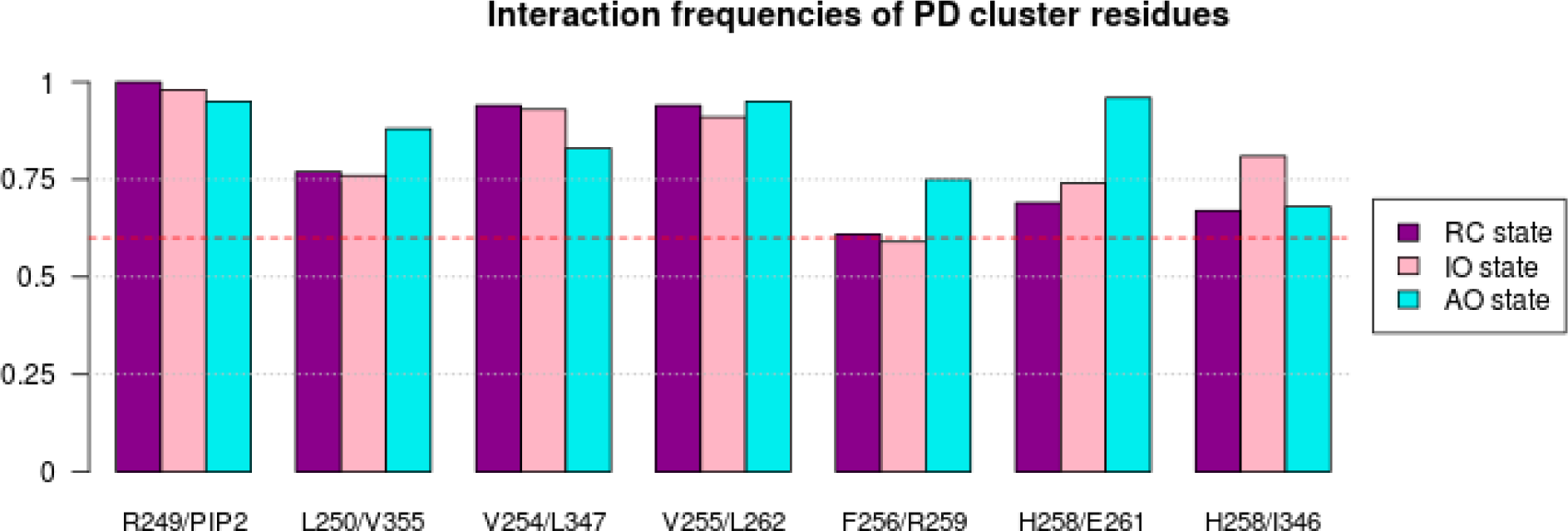
Intrasubunit interactions of PD cluster residues in K_V_7.1 models. The barplots display the frequencies of the state-independent intrasubunit interactions involving PD cluster residues throughout the MD trajectories obtained for K_V_7.1 RC (purple), IO (pink), and AO (cyan) models. The frequency threshold of 0.6 is marked by a red dashed line.

**Figure S3:**
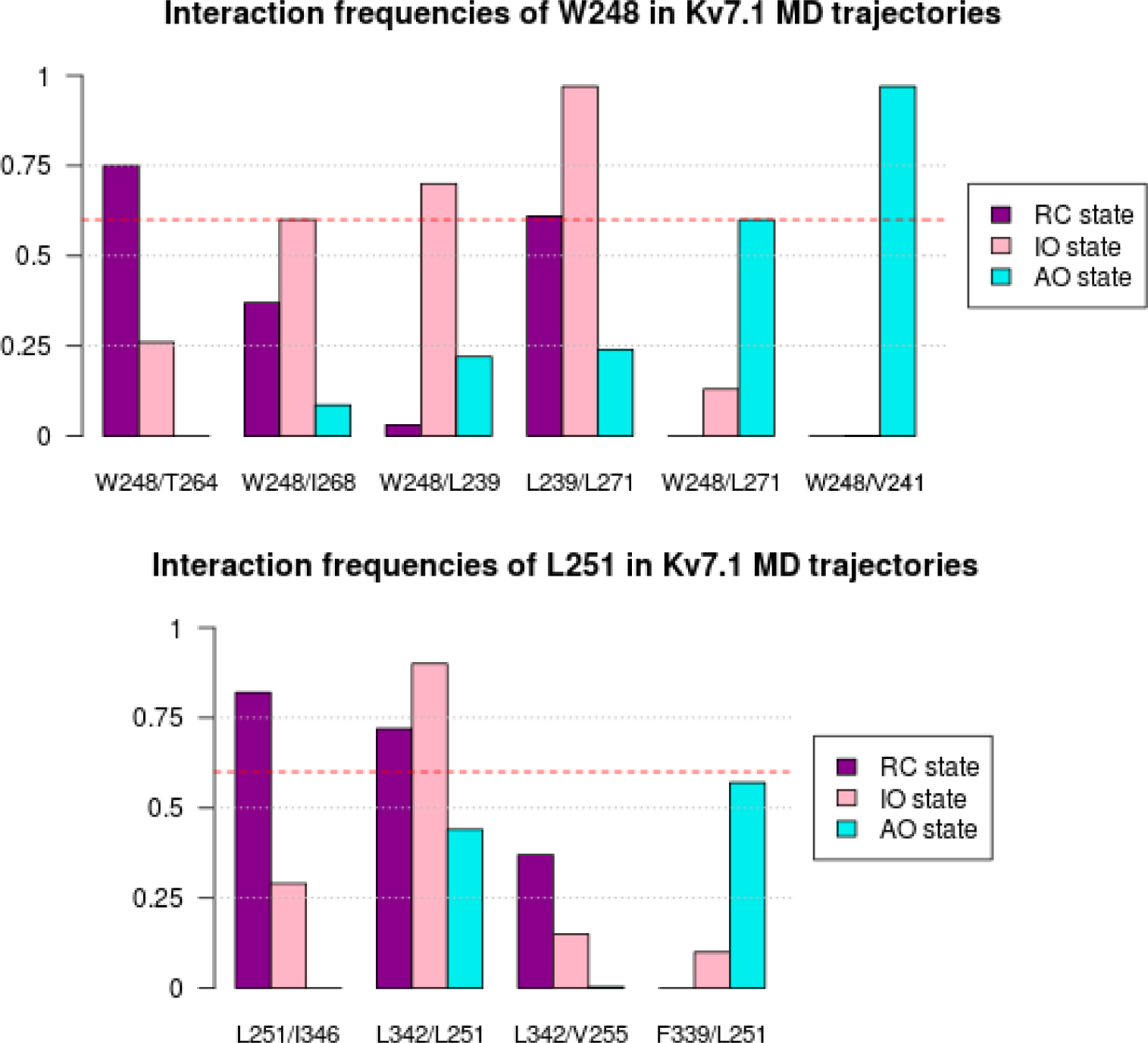
Intersubunit interactions of PD cluster residues in K_V_7.1 models. The barplots display the frequencies of the state-dependent intersubunit interactions involving PD cluster outer surface residues throughout the MD trajectories obtained for K_V_7.1 RC (purple), IO (pink), and AO (cyan) models. The frequency threshold of 0.6 is marked by a red dashed line.

**Figure S4:**
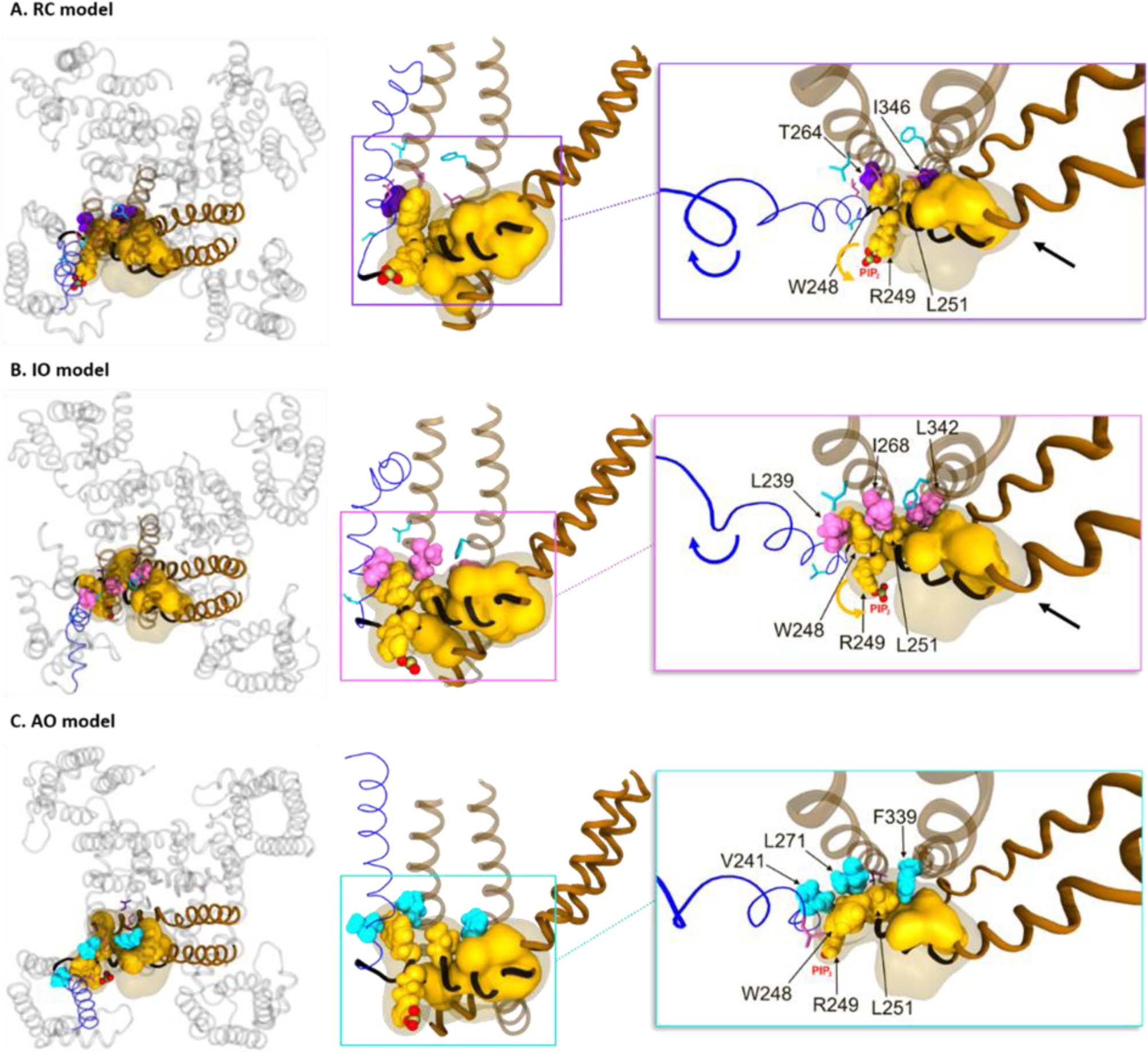
State dependent VSD-PD coupling interfaces in K_V_7.1 models. Each panel shows a cartoon representation of PD cluster residues in top view (left), lateral view (middle) and zoomed view (right) of **A.** RC model, **B.** IO model and **C.** AO model of K_V_7.1. The color code used for transmembrane segments is similar as in Figure 1,4 and S1. In all panels, the entire PD cluster is depicted in transparent yellow surface, and PIP2 phosphate groups in red spheres. Hook residues W248, R249, L251 and PD cluster inner surface residues are represented in solid yellow spheres and surface, respectively. In each panel, residues which interact with PD cluster are represented in spheres, while those which do not are represented in sticks. The residues which interact with PD cluster in **A.** RC models are purple-colored, while those which interact in **B.** IO model are pink-colored, and those which interact in **C.** AO models are cyan-colored. Rotational motions of S4, and PD cluster residues W248, R249, L251 are represented by blue, and yellow arrows, respectively. Translational movement of PD cluster is represented by a black arrow.

